# Control of G2 phase duration by CDC25B modulates the switch from direct to indirect neurogenesis in the neocortex

**DOI:** 10.1101/2021.12.14.472592

**Authors:** Mélanie Roussat, Thomas Jungas, Christophe Audouard, Francois Medevielle, Alice Davy, Fabienne Pituello, Sophie Bel-Vialar

## Abstract

During development, cortical neurons are produced in a temporally regulated sequence from apical progenitors, directly, or indirectly through the production of intermediate basal progenitors. The balance between these major progenitors’ types is determinant for the production of the proper number and types of neurons and it is thus important to decipher the cellular and molecular cues controlling this equilibrium. Here we address the role of a cell cycle regulator, the CDC25B phosphatase, in this process. We show that deleting CDC25B in apical progenitors leads to a transient increase of the production of TBR1+ neurons at the expense of TBR2+ basal progenitors in mouse neocortex. This phenotype is associated with lengthening of the G2 phase of the cell cycle, the total cell cycle length being unaffected. Using *in utero* electroporation and cortical slice cultures, we demonstrate that the defect in TBR2+ basal progenitor production requires interaction with CDK1 and is due to the G2 phase lengthening in CDC25B mutants. Altogether, this study identifies a new role for CDC25B and the length of the G2 phase in direct versus indirect neurogenesis at early stages of the cortical development.

## Introduction

During mammalian brain development, neural progenitor cells (NPCs) undergo precise sequential transitions that will define the number and type of neurons that will compose the neocortex (Kawaguchi 2019). It is therefore important that the proliferation and differentiation rates of these progenitors are tightly regulated during neurogenesis to avoid dramatic cortical defect such as microcephaly in human (Barbelanne and Tsang 2014).

At the onset of corticogenesis, NPCs initially divide symmetrically to expand the stem cell pool before giving rise to apical dividing progenitors (Govindan and Jabaudon 2017). Apical radial glial cells (aRGs) are located in the ventricular zone (VZ) and express Pax6. They first divide symmetrically and then shift to an asymmetric self-renewing division, producing an aRG and, either a neuron (direct neurogenesis) or a basal intermediate progenitor (bIP) expressing Tbr2. Basal progenitors divide once to produce two neurons (indirect neurogenesis) (Govindan and Jabaudon 2017, Kawaguchi 2019). Another type of basal progenitors, basal radial glia (bRG, also called outer radial glia), has emerged during evolution and is present in very small proportion in rodents in the SVZ and IZ. Like aRGs, they expresses Pax6 and have the capacity to perform several rounds of symmetric amplifying divisions (Ostrem, Di Lullo and Kriegstein 2017, Namba and Huttner 2017). All these progenitors sequentially produce the different types of projection neurons that will colonise the six layers of the adult neocortex (Kawaguchi 2019). The temporal transitions from one type of progenitor to another are therefore key parameters in controlling the size and functionality of the adult neocortex (Wilsch-Bräuninger, Florio and Huttner 2016).

Cell cycle kinetics and in particular cell cycle phase duration, is of major importance for controlling these transitions (Dehay and Kennedy 2007, Agius et al. 2015). Several studies showed that G1 lengthening is an important positive regulator of neuronal differentiation, G1-phase lengthening being associated with the transition from aRGs to bIPs during corticogenesis (Lange, Huttner and Calegari 2009, Pilaz et al. 2009, Arai et al. 2011). In addition it has been shown that proliferating and neurogenic progenitors display different S phase duration, the latter having a shorter S phase (Arai et al. 2011). Mitosis lengthening has also been linked to an increase in neurogenic divisions (Pilaz et al. 2016). However, even if correlations have been made in different organs between the duration of G2 phase and cell fate (Locker et al. 2006, Agathocleous et al. 2007, Gonzales and Liang 2015, Gonzales et al. 2015); the importance of controlling G2 length during neurogenesis remains poorly documented.

Few years ago, we have shown that the CDC25B phosphatase, a master regulator of mitosis entry, is expressed in neural progenitors and favours neurogenesis in the developing spinal cord (Benazeraf et al. 2006, Peco et al. 2012, Bonnet et al. 2018). CDC25B is a well-known G2/M regulator, belonging to the dual specificity CDC25 phosphatase family that activates CDK1-cyclinB complexes thus promoting mitosis entry (Boutros, Dozier and Ducommun 2006). It is also recruited to the mother centrosome and is involved in the centrosome duplication cycle as well as in microtubule nucleation (Boutros, Lobjois and Ducommun 2007, Boutros and Ducommun 2008, Boutros et al. 2011, Boutros et al. 2013). In the neural tube, CDC25B controls the G2 phase duration of neural progenitors, favours neurogenic division at the expenses of proliferative ones and, part of this function is independent of its interaction with CDK1 (Peco et al. 2012, Bonnet et al. 2018). CDC25B thus allows progenitors maturation during spinal neurogenesis. It remains an open question whether this phosphatase can be considered a general player in neuronal maturation and whether regulation of the duration of the G2 phase has a role to play in this process.

In this study, we investigated the role of CDC25B in corticogenesis. Using a conditional CDC25B knockout mouse line, we show that CDC25B loss of function in apical progenitors leads to a transient imbalance in neuronal and basal intermediate progenitors production, accompanied with a strong lengthening of the G2 phase. We provide evidence that this change in G2 duration in apical progenitors is causing the switch in fate, producing neurons instead of basal intermediate progenitors. This study sheds light on an important role of CDC25B and G2 phase modulation for controlling the fate of apical progenitors thus impacting the switch between direct to indirect neurogenesis.

## Results

### *Cdc25b* is expressed in apical progenitors

To determine if *Cdc25b* is expressed in all cortical progenitors or in a subset of them, we performed *in situ* hybridization on brain coronal sections at key stages of corticogenesis (**Figure 1A**).

**Figure 1:**
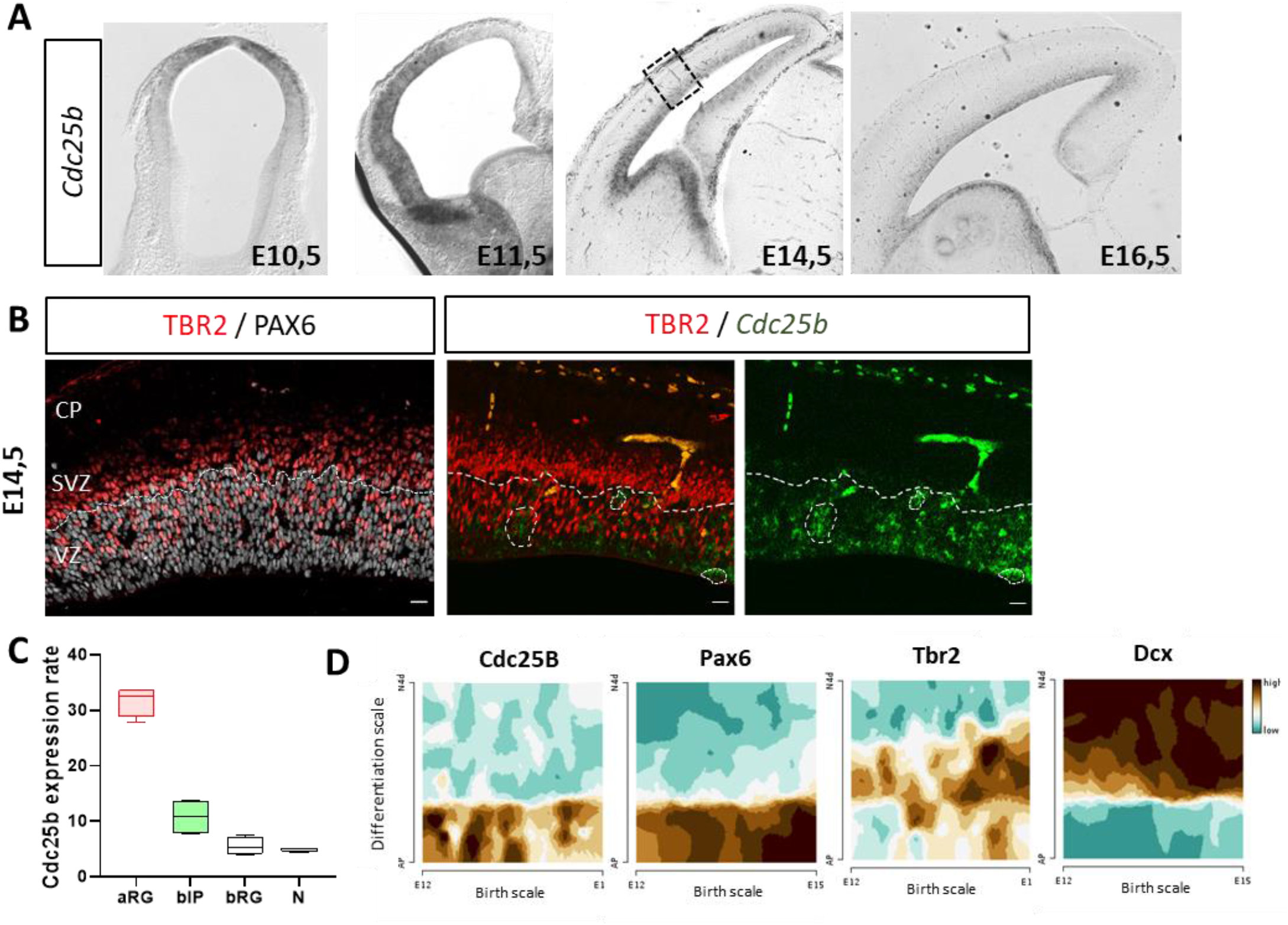
Cdc25b expression pattern during corticogenesis. Cdc25b is expressed in aRGs progenitors (PAX6+ in the VZ) and progressively downregulated in basal progenitors (TBR2+). **A,** Cdc25b in situ hybridization on mouse embryo coronal sections at E10.5, E11.5, E14.5 and E16.5. **B,** PAX6 and TBR2 immunostaining and co-staining of *Cdc25b* (in situ hybridization) and TBR2 expression (immunofluorescence) on E14.5 coronal brain sections. **C-D**, Quantification of CDC25B transcripts in progenitors and neurons using bulk RNA-Seq data from Florio et al.2015 (C) and Single-cell RNA-Seq from Telley et al 2019 using the gene browser at http://genebrowser.unige.ch/telagirdon/ (D).

*Cdc25b* is detected in neuroepithelial cells from E10.5. At E11.5, *Cdc25b* is expressed in the ventricular zone (VZ) in a graded manner: it is highly expressed laterally but at low level medially, its level of expression following the neuronal production gradient. At E14.5 and E16.5, it still appears restricted to the ventricular zone, that contains PAX6+ apical progenitors (aRGs) and TBR2+ newborn basal progenitors (bIPs) **(Figure 1A)**. To determine if it is also expressed in the subventricular zone (SVZ), we performed immunostaining for TBR2 following in situ hybridization of *Cdc25b* transcripts at E14.5. We observed that *Cdc25b* is not expressed in the SVZ, which contains TBR2+ bIPs, and that the TBR2+ cells in the VZ (newborn bIPs, Arai et al.2011) do not overlap with *Cdc25b*-expressing cells either **(Figure 1B)**. To confirm this observation, we analysed published bulk RNA-Seq data from E14.5 samples (Florio et al. 2015) and scRNA-Seq data from E12 to E15 cortices (Telley et al. 2019). This data mining confirmed that *Cdc25b* is highly expressed in aRGs, barely in bIPs and is not expressed in bRGs and neurons (**Figure 1C,D**). In addition, its expression appears to be most intense at early stages, when most bIPs cells are produced. Overall, these analyses indicate that in the developing cortex, *Cdc25b* is expressed gradually as neuronal production progresses and that it is predominantly expressed in aRGs, progressively extinguished in bIPs, and absent in neurons.

### *Cdc25b* deletion transiently leads to increased neuronal differentiation and decreased basal progenitor production

Having determined that *Cdc25b* expression is restricted to the apical progenitors, we next investigated its function during the early phases of corticogenesis in *Cdc25bfl/fl* ; *NestinCre (*CDC25B^cKO^ or cKO*) embryos* in which *Cdc25b* expression is specifically extinguished in neural progenitor cells from the onset of neurogenesis (Bonnet et al. 2018). In these embryos, deletion is complete in the cortex from E10.5 (Madisen et al. 2010) and at E14.5, *Cdc25b* expression is abolished in the whole brain **(Figure 2A)**. We first measured cortical hemispheres size and cortical plate thickness at postnatal stage (P0) and did not observed any gross difference in CDC25B^cKO^ embryos at these stages **(Figure 2B,C)**, suggesting that the number of neurons is unchanged. We then checked the production of different types of neurons over time. TBR1, CTIP2 and SATB2 expressing neurons are sequentially produced between E12.5 and E16.5 (Molyneaux et al. 2007, Vasistha et al. 2015). At E13.5, the majority of neurons express TBR1. In CDC25B^cKO^, the number of TBR1+ neurons is transiently increased in the most lateral part of the cortical plate at E13.5 **(Figure 2D,E)**. This increase is no longer significant at E14.5 **(Figure 2F,G)**. We then checked CTIP2+ and SATB2+ neurons at E14.5 and found that their production occurs normally at this stage in CDC25B^cKO^ **(Figure 2H,I,J)**. At E15.5 and E16.5, no difference is observed in the number of TBR1+, CTIP2+ or SATB2+ neurons (**Figure 2-figure supplement 1)**. These results indicate that in CDC25B^cKO^, neuronal production is transiently increased at E13.5 but this is compensated at later stages.

**Figure 2:**
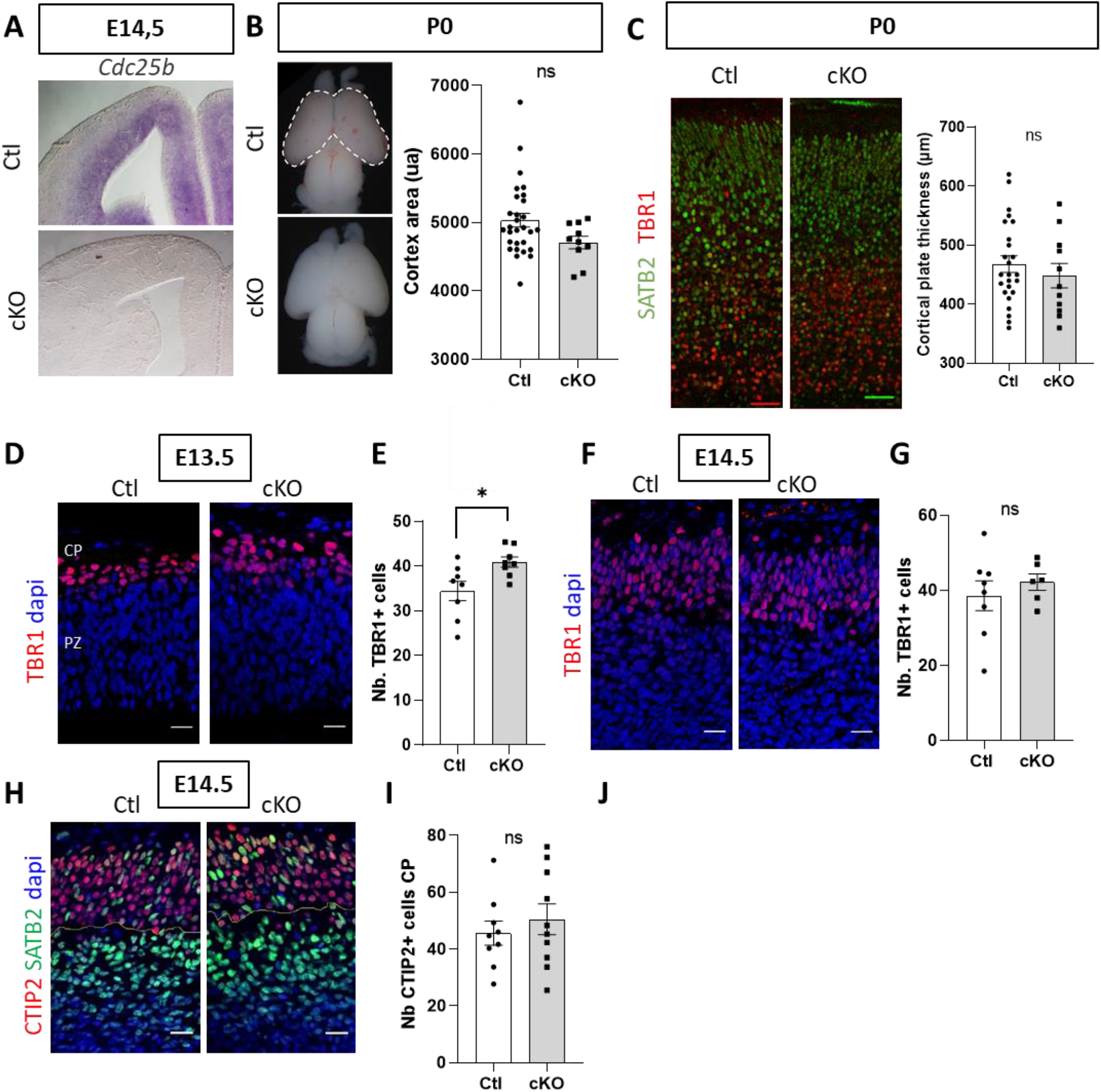
CDC25B loss-of-function triuersa transient increase of neuronal differentiation. **A**, Cdc25b in situ hybridization at E14.5 on control (Ctl) and Cdc25b cKO (cKO) embryo coronal sections. **B,** Cortical area (white dotted line) measurement in Ctl and cKO embryos at P0. Each point represents the cortex area in arbitrary unit (ua) for one embryo. Mann-Whitney t-test. Ctl n=29, cKO n=10 **C,** Cortical plate thickness measurement, delimited by SATB2^+^ and TBR1^+^ immunostaining on coronal brain sections of Ctl and cKO embryos at P0. Each point represents the value for one embryo. Mann-Whitney t-test. Ctl n=25, cKO n=11 **D, F**, TBR1 immunostaining on coronal sections at E13.5 (D) and E14.5 (F). Nuclei are stained with dapi (blue nuclei). **E, G**, Quantification of TBR1 neurons in the cortical plate (red nuclei) at E13.5 € and E14.5 (G). Each point is the mean value of 3 sections/embryo. Mann-Whitney t-test and mixed model. E13.5 n=9-10, *p<0.05; E14.5 n=6-8 **H**, CTIP2 (red nuclei) and SATB2 (green nuclei) immunostaining on coronal sections at E14.5, Nuclei are stained with dapi (blue nuclei). **I,J**, Quantificationof CTIP2 (I) and SATB2 neurons (J) in the cortical plate at E14.5. Each point is the mean value of 3 sections/embryo. Mann-Whitney t-test and mixed model. CTIP2 n= 9-10; SATB2 n=16. ns. Non significant. CP= cortical plate, PZ=progenitor zone, IZ=intermediate zone. Scale bars represent 20 μm.

Increased neuronal numbers at E13.5 could be due to an enhanced production of neurons from apical progenitors or to a premature differentiation of basal intermediary progenitors. To get insight on the dynamic of progenitor production in the CDC25B^cKO^ we examined aRGs distribution (PAX6+ cells in the VZ and SVZ), bIP cells production (TBR2+ newborn cells in the VZ) and bIP accumulation (TBR2+ cells that have migrated in the SVZ), between E12.5 and E16.5 **(Figure 3)**. The number of aRGs is not significantly affected at any of the 4 stages analyzed in CDC25B^cKO^ embryos (**Figure 3B,E,H,K**) and they are correctly located in the VZ (**Figure 3A,D,G,J**). At E14,5 the number of bIP is decreased in the VZ (**Figure 3G,I**). 48 hours later, the number of bIPs has returned to normal in the VZ but is now decreased in the SVZ (**Figure 3J,L)**. We also quantified PAX6+ cells in the IZ at E14.5 and E16.5 **(Figure 3M,N,O)** that are likely to be bRGS (Wang et al. 2011). The number of PAX6+ cells in the IZ is similar in Ctl and CDC25B^cKO^ at E14.5 **(Figure 3N)** but is increased at E16.5 **(Figure 3N,O)**.

**Figure 3:**
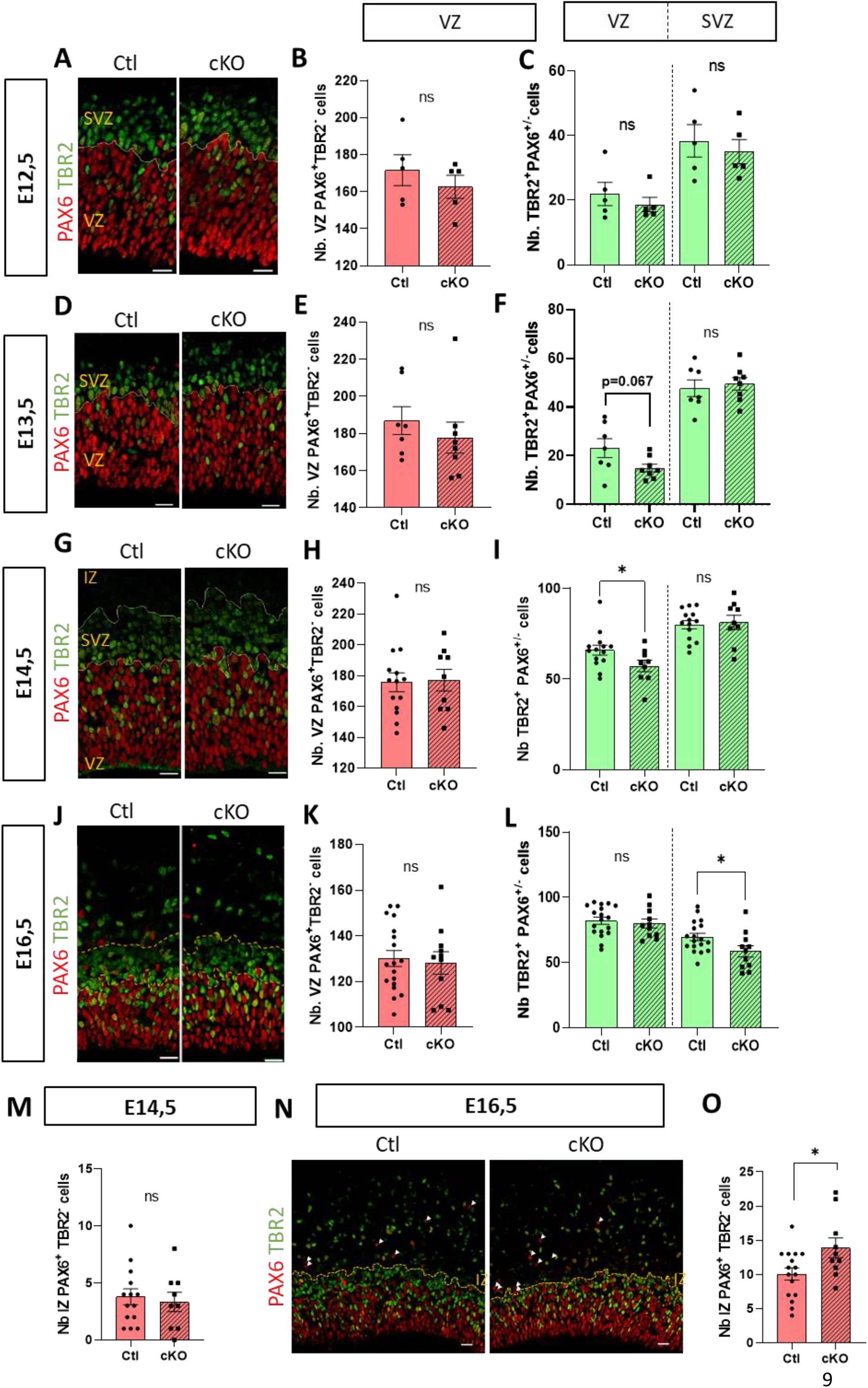
Cdc25b loss-of-function impairs basal progenitors production. **A, D, G, J, N** PAX6 and TBR2 immunostaining on E12.5 (A), E13,5 (D), E14,5 (G) and E16,5 (J) coronal brain sections, in Ctl and cKO embryos. **B-C, E-F, H-l, K-L**, Quantification of PAX6+ cells in the VZ (aRGs) and TBR2+ cells in the VZ and SVZ (bIPs) at E12.5 (B-C), n=5; E13.5 (E-F), n=7-8; E14.5 (H-I), n=9-14 or E16.5 (K-L), n=11-18 in Ctl and cKO embryos. Each point is the mean value of 3 sections/embryo. Mann-Whitney t-test and mixed model. **M, O**, Quantification of PAX6+ TBR2-cells in the intermediate zone (bRGs), at E14.5 (M), n=9-14 and E16.5 (O), n= 10-16 in Ctl and cKO embryos. Each point is the mean value of 3 sections/embryo. Mann-Whitney t-test and mixed model. * p<0.05, ** p<0.01, ns. Non significant. VZ = ventricular zone, SVZ = sub ventricular zone, IZ = intermediate zone. Scale bars represent 20 μ m.

To consolidate these data, we also quantified the total number of PAX6+ and TBR2+ cells in Control (Ctl) and CDC25B^cKO^ cortices using cytometry (Jungas et al. 2020). While the total number of PAX6+ cells is not modified, the number of TBR2+ cells is significantly reduced in CDC25B^cKO^ at E14.5 confirming that bIP but not aRGs production is affected in CDC25B^cKO^ **(Figure 3-figure supplement 1)**. We verified that this reduction was not due to apoptosis by quantifying activated Caspase3+ cells in the progenitor zone at E13.5 and E16.5 and, observed no difference between Ctl and CDC25B^cKO^ cortices. (**Figure 3-figure supplement 2A**). In addition, we quantified proliferation of bIPs (TBR2+ cells in VZ and SVZ) at E14.5 and found that proliferative and mitotic indexes were normal, ruling out that the reduced bIP number was due to a proliferation defect in these cells (**Figure 3-figure supplement 2 B,C,D**). Altogether, these results indicate that the reduced number of basal progenitors in CDC25B^cKO^ is most likely due to a reduction in their production from aRGs. This production defect is transient as the number of newborn TBR2+ cells in the VZ is not different from Ctl at E16.5. Altogether these results suggest that at E13.5, in absence of CDC25B, the progeny of aRGs is changed, from giving one aRG + one bIP to one aRG + one neuron. If this scenario is true, the balance between proliferative versus neurogenic division should not change in CDC25B^cKO^ as all three types of division mentioned above are neurogenic divisions.

To assess this, we generated CDC25B^cKO^ mice carrying a Tis21-GFP allele to distinguish aRGs committed to a neuronal or bIP fate (TIS21+) from self-expanding aRGs (TIS21-) (**Figure 3-figure supplement 3)** (Haubensak et al. 2004, Attardo et al. 2008). We did not detect any change in the number of TIS21-GFP cells among aRGs at E13.5 in CDC25B^cKO^ compared to Ctl (**Figure 3-figure supplement 3**), confirming that CDC25B does not change the mode of division of aRGs from proliferative to neurogenic but rather influence the fate of neurogenic divisions. Overall, these analyses of neuron and progenitor production at several developmental stages indicate a transient increase in neuron production accompanied by a transient decrease in bIP production in CDC25B^cKO^ embryos, with no change in aRG numbers. This suggests that in CDC25B^cKO^, aRGs transiently produce neurons instead of bIPS and that CDC25B controls the switch between direct versus indirect neurogenesis.

### The control of basal progenitor production by CDC25B requires CDK interaction

We previously showed that CDC25B can act independently of CDK interaction and G2 phase regulation in the developing spinal cord (Bonnet et al. 2018). To test if its role in cortical neurogenesis was dependent or independent of CDK interaction, we compared the effect of CDC25B gain-of-function with that of a mutated form of CDC25B, CDC25BdelCDK, which is no longer able to interact with CDK1 to regulate G2 phase length (Bonnet et al. 2018). We used *in utero* electroporation at E13.5 and analyzed the fate of electroporated cells 28h later **(Figure 4A)**. Compared to a GFP+ control vector, ectopically expressing CDC25B increases the number of TBR2+ cells without changing the number of aRGs (TBR2-cells in the VZ), indicating that CDC25B is sufficient to promote bIPs production **(Figure 4B-D)**. Unlike WT CDC25B, ectopic expression of CDC25BdelCDK did not alter the number of TBR2+ cells **(Figure 4B-C)**. These results clearly show that CDC25B is sufficient to enhance bIPs production and that this effect requires an interaction with CDK. This suggests that this control is exerted via a modification of the duration of the G2 phase.

**Figure 4:**
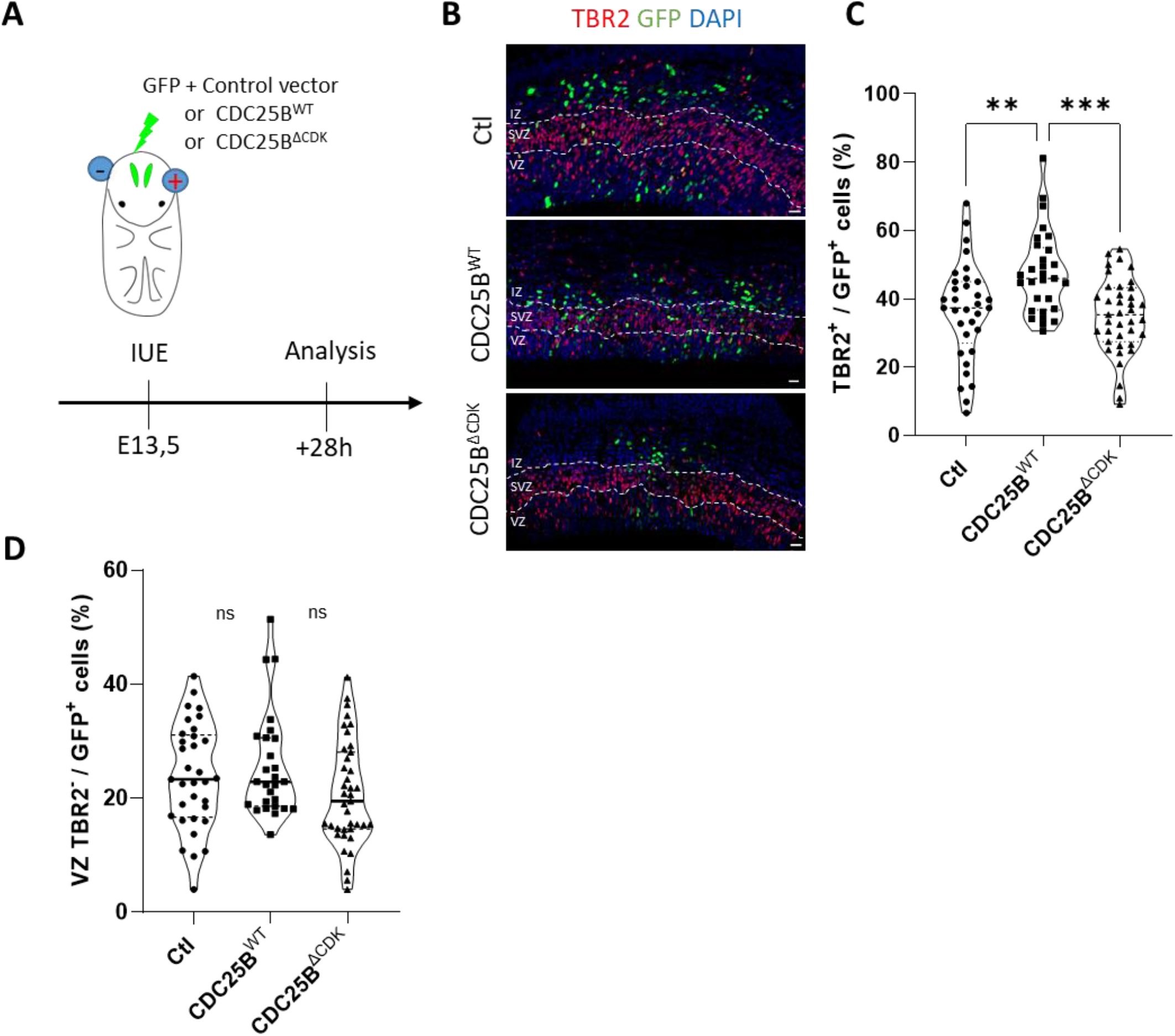
Cdc25b regulation of bIP production is a CDK1-dependant function. **A**, Schematic representation and timeline of in utero electroporation experiments. **B,** GFP and TBR2 immunostaining on E14.5 brain coronal sections after in utero electroporation of a pmp-LacZ (CTL), pmp-CDC25BWT (CDC25BWT) or pmp-CDC25BdelCDK (CDC25BΔCDK) expression vectors at E13.5. **C,** Quantification of GFP+ cells which are TBR2+ (bIPs) per section of CTL, CDC25BWT or CDC25BΔCDK electroporated embryos. **D,** Quantification of GFP+ cells that are TBR2-in the ventricular zone (aRGs) after electroporation. Each point represents the mean value for 3 z-section/embryo slice. Kruskal-Wallis test. Ctl n=29, cKO n=39, **p<0.01 ***p<0.0001. ns. Non significant. VZ = ventricular zone, SVZ = sub ventricular zone, IZ = intermediate zone. Scale bars represent 20 μ m.

### Cdc25b loss-of-function severely increases the length of the G2-phase without affecting the total duration of the cell cycle

Knowing the role of CDC25B in the control of G2/M cell cycle phase transition, and as bIP production involves CDC25B/CDK interaction, the defect in bIP production could be due to a defect in cell cycle parameters in aRGs of CDC25B^cKO^ embryos. We first measured the duration of the G2 phase using the percentage of labeled mitoses (PLM) paradigm (Quastler and Sherman 1959). Embryos were harvested 2 hrs, 3 hrs, 4 hrs (**Figure 5A**) or 6 hrs (not shown) after EdU (5-ethynyl-2′-deoxyuridine) administration and we quantified the percentage of P-H3^+^(Phospho-histone 3) EdU^+^/P-H3+ with increasing exposure times to EdU (**Figure 5A,B**). We found that this percentage was consistently lower in aRGs from CDC25B^cKO^ embryos.

**Figure 5:**
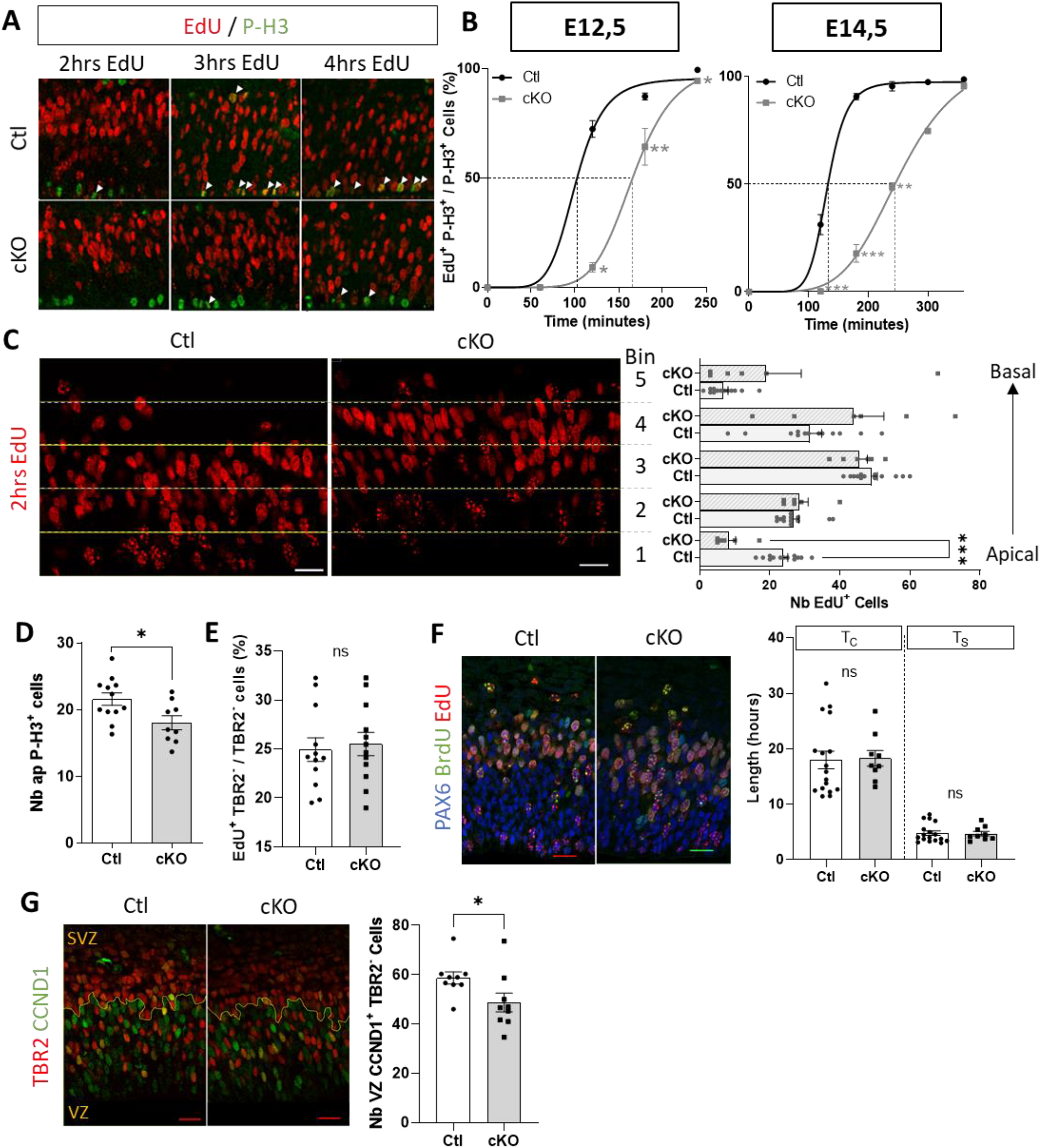
Cdc25b removal strongly lengthens the G2-phase of aRG progenitors. **A**, PH3 labelling after 2, 3 and 4 hours of EdU pulse in Ctl and cKO embryos at E14.5. **B,** Quantification of the proportion of apical (aRG) EdU^+^ P-H3^+^ /P-H3^+^ cells (A, arrows). T_G2_ corresponds to the time when 50% of P-H3^+^ cells are EdU (indicated by the dotted lines). Mann-Whitney t-test. N=2-10, *p<0.05, **p<0.01, ***p<0.0001 **C,** EdU^+^ cell quantification after 2 hours EdU pulse, in 30 μm high bin from apical (bin 1) to basal side (bin 5), in Ctl and cKO embryos at E14.5. Mann-Whitney t-test. N=6-13, ***p<0.0001 **D,** Quantification of apical P-H3+ cells in Ctl and cKO E14.5 cortices. Mann whitney and Mixed model. n=9-12, *p<0.05 **E,** Quantification of EdU^+^ cells among VZ TBR2^−^ (aRGs) cells after 1 hour EdU pulse in Ctl and cKO E14.5 embryos. Mixed model. n=12 **F,** EdU/BrdU double labelling experiments to determine total cell cycle length (T_c_) and S-phase duration (T_s_) of VZ PAX6^+^ cells (aRG), which were calculated from counts of EdU^+^BrdU^−^ and EdU^+^BrdU^+^ cells as described (Martynoga et al., 2005). Bar plots represent the average length of measured T_c_ and T_s_ in aRGs of Ctl and cKO E14.5 embryos. Mann whitney and Mixed model. n=9-17 **G,** CCND1/TBR2 double immunostaining to determine the proportion of progenitor in Gl-phase. Quantification of CCND1^+^ cells among VZ TBR2^−^ (aRGs) cells in Ctl and cKO E14.5 cortices. Mixed model. n=9, *p<0.05. VZ = ventricular zone, SVZ = sub ventricular zone. In bar plots, each point is the mean value of 3 sections/embryo. Scale bars represent 20 μ m.

Because the progression of this percentage is proportional to the duration of the G2 phase, we were able to extract an average length of the G2 phase. Loss-of-function of *Cdc25b* in aRGs lead to a significant increase in the duration of the G2 phase at E14.5, from 2h12 hours in controls to 4h05 in CDC25B^cKO^ (**Figure 5B, Table 1)**. This lengthening is already detected at E12.5 and lasts at least until E16.5 **(Figure 5B –figure supplement 1)**. In addition, we observed a strong delay in apical migration of EdU-labeled cycling progenitors **(Figure 5C)**, linked to a reduction in the number of apical mitosis (P-H3 positive cells, **Figure 5D**), coherent with the G2 phase lengthening. We then determined whether this lengthening of G2 phase duration in progenitors affects their proliferation rate. We collected embryos after 1h of EdU administration at E14.5 and quantified the EdU+ cells among the aRGs (VZ, TBR2-cells, **Figure 5E**). This percentage reflects the proportion of aRGs in S phase and we did not observe any difference between Ctl and CDC25B^cKO^ embryos (**Figure 5E)**. As the rate of proliferating cell was unaffected, we next investigated whether the lengthening of the G2 phase caused a general slowdown of the total cell cycle length in aRGs. We performed dual EdU and BrdU (bromodeoxyuridine) labelling to estimate total cell cycle length (Tc) and S phase (TS) durations, as described in (Martynoga et al. 2005). As shown in (**Figure 5F)**, nor the total cell cycle length neither TS phase are significantly modified in aRGs of CDC25B^cKO^ embryos. From this result, we conclude that a shortening of the G1 phase duration may compensate for the strong lengthening of the G2 phase. To confirm this hypothesis, we directly assessed the length of the G1 phase by counting the number of cells expressing CCND1 (specifically expressed during the G1 phase) among the aRGs. We observed a reduction in the number of CCND1+ labelled nuclei in CDC25B^cKO^ aRGs (**Figure 5G**) indicating that the G1 phase is shorter in these cells.

**Table 1:** Cell cycle parameters in Ctl and cKO aRGs

| E14,5 | T<sub>C</sub> | T<sub>S</sub> | T<sub>G2</sub> | *T<sub>G1</sub> |
| --- | --- | --- | --- | --- |
| Control | 17h58 | 4h47 | 2h12 | 10h29 |
| Cdc25b<sup>nesKO</sup> | 18h15 | 4h41 | 4h05 | 8h59 |
| $\Delta T$ | + 0h18 | - 0h06 | + 1h53 | - 1h30 |
| % FC | + 1,7% | - 2% | + 85% | - 14% |

In conclusion, the removal of CDC25B in aRGs leads to a strong lengthening of the G2 phase (85% length increase, **Table 1**) which is compensated by a shortening of the G1, resulting in a normal cell cycle length and proliferation rate. If these length changes are considered in proportion to their average length, the strongest change we observed is in the duration of the G2 phase **(Table 1)**.

### G2-phase shortening drives an increase in bIP production

As the duration of the G2 phase is almost doubled in CDC25BcKO, we next investigated whether this lengthening could play a role in controlling the rate of bIP production. To this end, we turned to cortical slice cultures and applied pharmacological treatments to modulate the duration of the G2 phase. First, we treated cortical slices of E13.5 WT embryos with PD0166285, a Wee1/Myt1 inhibitor in conditions that do not induce toxicity **(Figure 6A)**(Baffet et al.2015). As Wee1 and Myt1 prevent mitosis entry through negatively regulating CDK1 activity (Potapova et al. 2009), their inhibition leads to an increase in CDK1 activity and thus to a shortening of the G2 phase. We first quantified that PD0166285 treatment at 1um has an effect on G2 length in our cultures. For that, we quantified the number of EdU+/P-H3+ cells among the P-H3+ cells following 2h EdU incorporation just before culture arrest. The percentage of EdU-positive cells is identical between untreated and treated controls (approximately 22% in both conditions, not shown) indicating that the rate of proliferation is not significantly altered by pharmacological treatment and that the cortical slices are still in a good physiological state at the end of the culture. We confirmed that this treatment effectively reduces the time spent in G2 phase by considering the percentage of EDU+P-H3+ /P-H3+ cells in untreated and treated samples **(Figure 6B,C)**. We analyzed the impact of this treatment on the duration of the G1 and S phases and on the total duration of the cell cycle. We found that S-phase is significantly reduced by PD0166285 treatment, while G1-phase duration and total cell cycle length are not significantly modified **(Figure 6D,E)**. We then counted the number of TBR2+ cells in treated and untreated slices and observed that the pharmacological shortening of the G2 phase is accompanied by an increase in the number of bIPs in the VZ **(Figure 6F,G-supplement figure 1)**. This indicates that PD0166285 treatment mimics the effect of CDC25B gain of function on number of TBR2+ progenitors. We then investigated whether this treatment would be sufficient to restore normal numbers of TBR2+ cells in CDC25B^cKO^ cortices. We therefore applied the same experimental protocol as above to CDC25B^cKO^ and control littermates. As observed on WT embryos, treatment with 1uM PD0166285 led to a marked increase in EdU+P-H3+/P-H3+ cells in both control and CDC25B^cKO^ -treated slices, indicating a shortening of the G2 phase, with the EDU+P-H3+/P-H3+ ratio in treated CDC25B^cKO^ slices now at the level of untreated Ctl littermates **(Figure 6H)**. We then checked the number of TBR2+ cells in the VZ of these treated and untreated embryos and observed a significant increase in these cells in CDC25B^cKO^ and Ctl littermates-treated slices **(Figure 6I,J)**. This indicates that shortening the G2 length in CDC25B^cKO^ is sufficient to restore efficient bIPs production.

**Figure 6:**
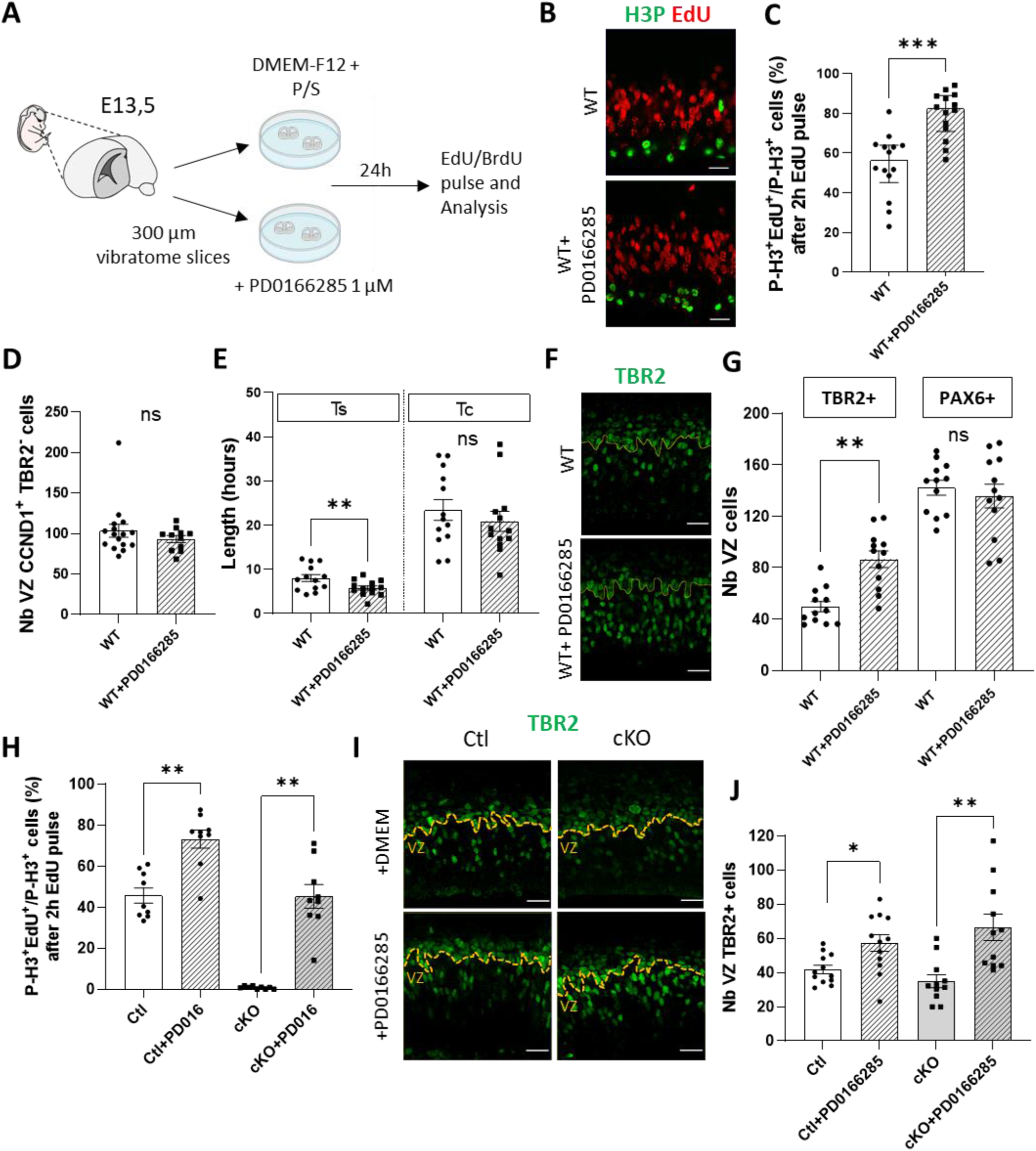
G2-phase shortening triggers bIP production. **A**, Schematic representation of embryo brain slice culture experiments. **B,** P-H3 labelling after 2h of EdU pulse in Ctl and cKO embryos at the end of the culture **C,** Percentage of EDU^+^H3P^+^/H3P^+^ cells following 2 h EdU pulse, in untreated and PD0166285 treated WT brain slices. Wilcoxon test. n=14,***p<0.0001 **D,** Quantification of CCND1^+^ TBR2^-^ cells (aRG in Gl-phase) in WT and WT+PD0166285 brain slices. Wilcoxon test. n=11-16 **E,** Total cell cycle length (T_c_) and S-phase duration (T_s_) of VZ TBR2^-^ cells (aRGs) in WT and WT+PD0166285 brain slices. Wilcoxon test. n=13, **p<0.01 **F,** TBR2 immunostaining on WT and WT+PD0166285 brain slices **G,** Quantification of bIPs (TBR2+ cells) and aRGs (TBR2-cells) in the VZ of WT and WT+PD0166285 brain slices. Wilcoxon test. n=17, **p<0.01 **H,** Percentage of EDU^+^H3P^+^/H3P^+^ cells following a 2h EdU pulse in untreated or PD0166285 treated, Ctl or cKO brain slices. Wilcoxon test n=9, **p<0.01 **I,** TBR2 immunostaining on untreated or PD0166285 treated Ctl or cKO brain slices **J,** Quantification of TBR2+ cells (bIPs) in the VZ of untreated or PD0166285 treated, Ctl or cKO brain slices. Wilcoxon test. n=9, *p<0.05, **p<0.01. Each point represents the mean value of 3 sections/embryo. VZ = ventricular zone. Scale bars represent 20 μm

Overall, these results indicate that the duration of the G2 phase is an important parameter to control bIP production during early phases of cortical development.

## Discussion

In this study, we investigated the function of CDC25B during corticogenesis. We show that removing CDC25B function lead to a transient increase in neuron numbers at early stages, accompanied with a decrease in the number of intermediate basal progenitors. This imbalance of neuron/progenitors number is due to a modification of the G2 phase length in apical progenitors. We propose a model in which CDC25B expression in aRGs, controls the switch from direct to indirect neurogenesis through a modulation of G2 phase length.

In the developing neocortex, neurons can be produced either directly from asymmetric division of apical progenitors or indirectly from symmetric division of intermediate basal progenitors (Govindan & Jabaudon, 2017). Controlling the balance between these two modes of production is important for the integrity and size of the cortex, but also has an impact on the type of neurons that are produced, (Haubensak et al., 2004, Attardo et al., 2008, Govindan & Jabaudon, 2017). In CDC25B^cKO^, we observed a specific increase of TBR1+ neurons at E13.5 followed by a decrease in bIPs numbers at E14.5. Conversely, a gain of function of CDC25B at E13.5 leads to an increase in bIPs in the VZ. We can exclude that the increase in neuron production is due to an increase in aRG production because the number of aRG is not altered in CDC25B^cKO^. Furthermore, if this scenario were true, one would expect that there would also be more bIPs produced. This alteration in the number of bIPs and neurons is also not due to a reduction of neuron production from bIPs. Indeed, the increase in neuron production in CDC25B^cKO^ embryos precedes the decrease in bIPs, and if bIPs differentiated into neurons prematurely, we should detect a decrease in bIPs cells in the SVZ as early as E13.5, which is not the case. Since the number of aRGs is normal and the ratio of proliferative to neurogenic division is not changed in CDC25B^cKO^, our results indicate a change in the progeny of asymmetrically dividing aRGs, producing neurons instead of bIPs (**Figure 7**). This phenotype is transient, as at E16.5 we no longer observe a defect in bIP numbers in the VZ and, the number of neurons is unchanged in CDC25B^cKO^. This could indicate either a compensation mechanism (Betty et al. 2016) or alternatively, that CDC25B is only required at early corticogenesis stages, at the time of the direct to indirect neurogenesis switch. Hence, we propose that in CDC25B^cKO^, the switch from direct to indirect neurogenesis is delayed, leading to a bIP/neuron production imbalance which is progressively compensated (**Figure 7**).

**Figure 7:**
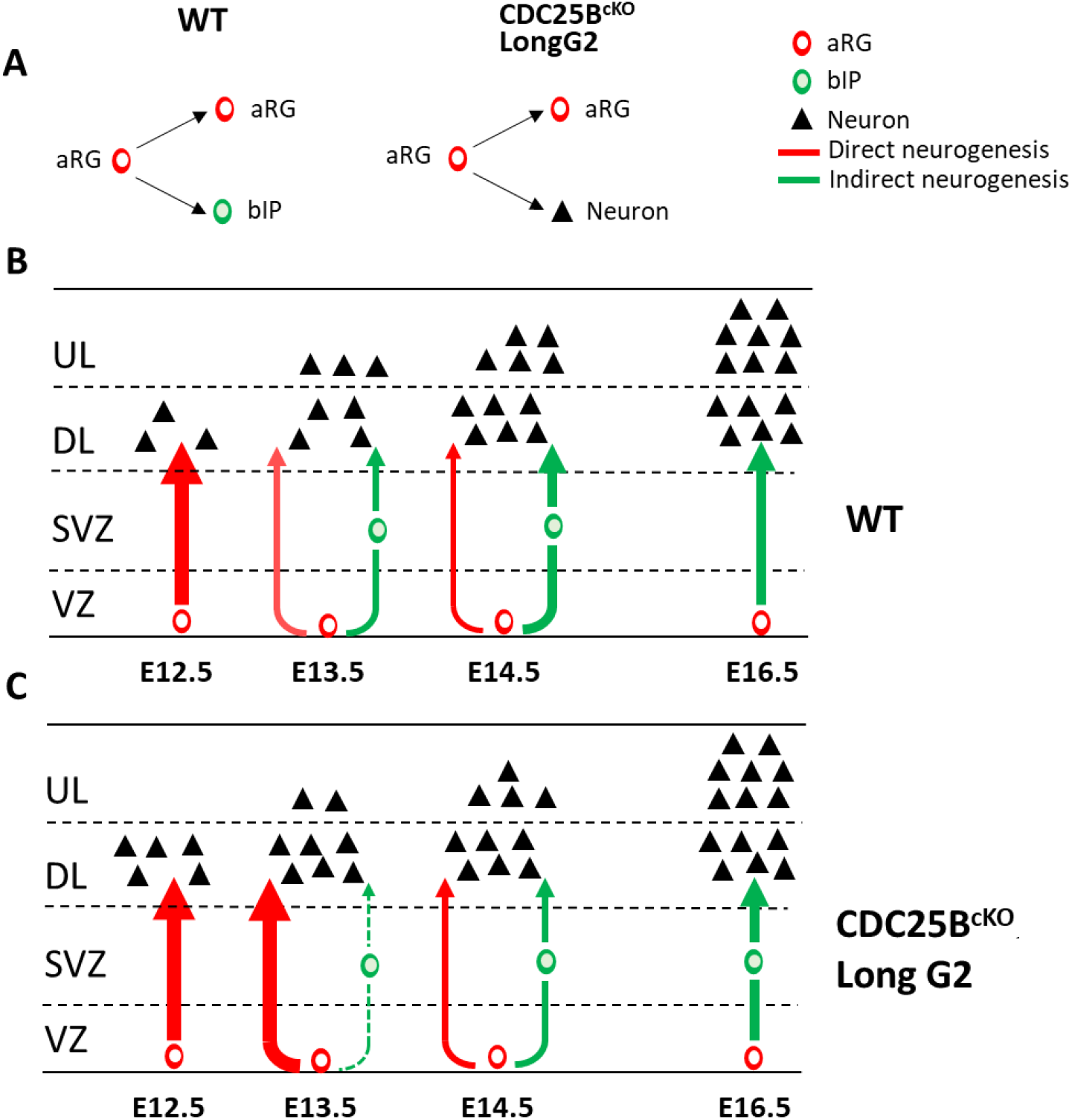
Model for CDC25B function during cortical neurogenesis. **A**, In CDC25B^cKO^ apical progenitors asymmetric division produce a neuron instead of a basal progenitor **B**, In a normal situation, deep layer neurons are produced mainly between E12.5 and E13.5, directly from APs (direct neurogenesis). At E13.5, the production of bIPs begins in the VZ, which later divide in the SVZ to give rise to neurons (indirect neurogenesis). Indirect neurogenesis gradually becomes the main mechanism of neuron production. **C**, In CDC25B^cKO^, the ratio between direct and indirect neurogenesis is disturbed. Direct neurogenesis is maintained for a longer period of time, leading to an early production of deep layer neurons at the expense of the production of bIPs. This imbalance is progressively compensated to reach normal neuronal production around E16.5.

In a previous study, we have shown that CDC25B promotes neurogenesis in the developing spinal cord by increasing neurogenic divisions at the expense of proliferative ones (Bonnet et al. 2018). Other studies have also made indirect links between defects in the balance between proliferative versus neurogenic division during cortical neurogenesis and CDC25B misregulation (Gruber et al. 2011, Wu et al. 2014). We did not see any change in the proliferative versus neurogenic mode of division in CDC25B mutant cortices. Our previous observations made in the chicken neural tube, indicate that part of CDC25B function during neurogenesis is independent of its function on the cell cycle (Bonnet et al. 2018). In the neocortex, the phenotype we describe requires interaction with CDK1, but in both tissues CDC25B appears to promote progenitor maturation.

Another important result is that this neurogenesis defect is linked to a strong modification of the G2 phase length. First, in CDC25B^cKO^, the most important change in cell cycle parameters in aRGs is a strong lengthening of the G2 length. Second, the gain of function of CDC25BdelCDK, that cannot regulate G2 phase length (Bonnet et al. 2018) does not phenocopy the wild-type form of CDC25B that increases bIPs production, indicating that the defect in bIPs production is directly related to the ability of CDC25B to regulate CDK1 activity. Third, we provide evidence that shortening the G2 phase in WT embryos using a pharmacological treatment is sufficient to stimulate bIP production and fourth, that this treatment is sufficient to restore an efficient bIP production in CDC25B^cKO^. In these experiments, either the G1 phase length or the S-phase length were slightly reduced. However, the levels of change in these phases are well below the levels that have been described to result in changes in aRGS progenitor mode of division and fate (Lange et al. 2009, Pilaz et al. 2009, Arai et al. 2011). Furthermore, the only parameter that varies strongly and significantly in both CDC25B^cKO^ and PD0166285 treatment is the length of the G2 phase. This argues for a role for the duration of the G2 phase rather than for a role for the G1 or S phase durations in these experimental settings. In CDC25B^cKO^, the G2 lengthening is associated with a perturbation of the interkinetic nuclear movement as BrdU+ aRGs nuclei take more time to reach the apical surface to perform mitosis. This perturbation is likely due to the fact that INM and G2 phase processing are linked and, cell cycle progression is a prerequisite to normal INM (Ueno et al. 2006, Baffet, Hu and Vallee 2015). At later stages (E16.5), the duration of the G2 phase is still greatly extended in CDC25B^cKO^ but neuronal numbers have been compensated suggesting that G2 length modulation is important specifically at the time of the switch between direct versus indirect neurogenesis but not for later steps of neuronal production.

What is the relevance of this finding on the link between G2 phase length and bIP production in a normal context? Most studies in which GAP phase length has been measured indicate that while G1 phase length increases during tissue maturation, G2 phase length appears to be more stable, around 2h in aRGs and bIPs and, does not change significantly during embryonic cortex maturation (Pilaz et al. 2009, Arai et al. 2011, Hasenpusch-Theil et al. 2018). However, these studies were performed at the level of the cell population, which could have masked some heterogeneity in the duration of the G2 phase between different progenitors. Data provided using real time analysis of PCNA expressing apical progenitors show that the proportion of G2 phase in aRGs change between E12 and E16 (Fousse et al. 2019). A recent study from Hagey et al. also emphases that cortical progenitors are not equivalent regarding G2/M phase length (Hagey et al. 2020). Indeed, by performing single cell RNA seq from E9.5 to E18.5, they show that at E11.5, progenitors can be separated in two groups, one of committed progenitors, giving rise to deep layer neurons, the other one of uncommitted progenitors that will give rise to later born lineages. Interestingly, the group of committed progenitors spend more time in G1 and express G1 associated cyclin, whereas the uncommitted ones are more prone to be found in G2/M and express high levels of M-phase associated genes, including CDK1. Interestingly, data mining of this study and others (Telley et al. 2019, Hagey et al. 2020) indicate that *Cdc25B* transcripts have the same spatial and temporal profile than genes specifically expressed in the uncommitted E11.5 progenitors. Hence, CDC25B activity, by accelerating mitosis entry, could contribute to the maintenance of the uncommitted progenitor pool. Different G2 length duration, by changing the duration of exposure to instructive signaling pathways such as the Slit-Robo and/or the Notch signaling pathways (Murciano et al. 2002, Cárdenas et al. 2018, Fousse et al. 2019, Hagey et al. 2020) could lead to different fate decision after mitosis.

In conclusion, our study shed light on a new role for CDC25B and G2 phase length in controlling the fate of apical progenitors during early steps of corticogenesis, thus controlling the switch between direct and indirect neuronal differentiation. The apparition of indirect neurogenesis is considered a milestone of mammalian cortical evolution leading to cortical expansion (Borrell 2019). Expression of CDC25B in aRGs, by favoring bIP production, would be a way of supporting this evolutive trait. Future investigations will be necessary to understand the molecular mechanisms that support this function.

## MATERIAL AND METHODS

### Mice

Experiments were performed in accordance with European Community guidelines regarding care and use of animals, agreement from the Ministère de l’Enseignement Supérieur et de la Recherche number: C3155511, reference 01024.01 and CNRS recommendations. Cdc25b +/- ; Nestin-Cre, and Cdc25b flox/flox lines has been described previously (Tronche et al. 1999) (Bonnet et al. 2018). These lines were crossed to obtain conditional mutant embryo Cdc25 flox/-; nestin-Cre (cKO) and control embryos Cdc25b flox/-, Cdc25b flox/+ and Cdc25b flox/+; nestin-Cre (Control). Tis21-GFP line *(Tis21*^*-/tm2(Gfp)Wb*^) has been described previously (Haubensak et al., 2004). This line was crossed to obtain conditional mutant embryo Cdc25b flox/-; nestin-Cre; Tis21-GFP, that express GFP under the control of Tis21 promotor. Embryonic (E) day 0.5 was assumed to start at midday of the day of the vaginal plug.

### In situ hybridization and immunohistochemistry

In situ hybridization were performed on 60μm vibratome sections of mouse brain embryos as described (Bonnet et al. 2018). For immunohistochemistry, embryos were fixed in 4% paraformaldehyde at 4°C overnight, washed in PBS and embedded in paraffin using an STP 120 Spin Tissue Processor. Brains were sliced in 6 to 14 μm transversal sections using a HM 355 Microtom. Section are directly mounted on coated histological slides. Sections were rehydrated by successive baths of histoclear (2 times), 100% ethanol (2 times), 95% ethanol, 70% ethanol, 50% ethanol and deionized H_2_0 for 5 minutes each. Sections were heated in 1% Citrate in H_2_O at around 100°C for 45 minutes, permeabilized with PBS-0,5% Tween + 3% Bovine Serum Albumin + 1% Horse Serum and incubated with primary antibody overnight at 4°C. The following primary antibodies were used against: P-H3 (Upstate Biotechnology, 1:1000, rabbit), BrdU (G3G4, 1:400, mouse), active caspase 3 (BD Biosciences, 1:100, rabbit), GFP (abcam, 1:1000, chick), CCND1 (ThermoFisher, 1:200, rabbit), SATB2 (Abcam, 1:100, mouse), TBR1 (abcam, 1:100, rabbit), PAX6 (Covance, 1:300, rabbit), PAX6 (MBL; 1:200, rabbit), TBR2 (eBioscience, 1:50, rat), RFP (SICGEN, 1:100, goat), γ-Tubulin (Exbio, 1:1000, mouse), CTIP2 (abcam, 1:200, rat). Sections were then washed in PBS and incubated with the secondary antibodies for 2h at room temperature. We counterstained the sections with DAPI (1:1000, Life Technologies) to identify nuclei. For BrdU detection, we treated sections with 2N HCl and then with 0.1 M Na_4_B_4_O_7_ before incubation with the primary antibodies.

### Cell proliferation, cell cycle and birth dating analysis / EdU/BrdU experiments

We determined G2-phase length using the percentage of labeled mitoses (PLM) paradigm (Quastler and Sherman 1959). We injected pregnant mice intraperitoneally with 100 μL of 5-ethynyl-2’-deoxyuridine (EdU, Click-iT EdU Alexa Fluor 647 Imaging Kit, Invitrogen) at 1mg/mL. Embryos were harvested 120, 180, 240, 300, 360 or 480 minutes after EdU administration and, we quantified the percentage of P-H3^+^EdU^+^ co-labeled nuclei with increasing times of exposure to EdU. The progression of this percentage is proportional to G2-phase duration. To determine cell proliferation, we injected pregnant mice intraperitoneally with 1 mg/mL of EdU. Embryos were harvested 60 minutes later. Total cell cycle length and S-phase length were determined as described (Martynoga et al., 2005). We injected pregnant mice intraperitoneally with 100 μL of EdU 10 mg/mL, followed by 100μL of 5-bromo-2’-désoxyuridine (BrdU, Sigma) at 10 mg/mL 90 minutes later. Embryos were collected 30 minutes later. We calculated Tc and Ts from counts of EdU^+^BrdU^−^ and EdU^+^BrdU^+^ cells. To determine the generation of neurons by pulse-chase experiments (birth dating), we injected pregnant mice intraperitoneally with EdU at E16,5 and we harvested embryos at birth (P0). We deduced the G1 phase length by removing the average length of G2+S+M phases from the average total cell cycle length in control and CDC25B^cKO^ littermates.

### Flow cytometry analysis

Flow cytometry analysis were conducted as described in (Jungas et al. 2020). Cortical hemispheres were collected in PBS-1% FCS and cells mechanically dissociated through a 40-μm nylon mesh (Clearline). We suspended cells in 1.2 ml ice-cold PBS and fixed them by slowly adding 3 ml cold 100% ethanol while vortexing to obtain a 70% final concentration. The incubation procedure with primary and secondary antibodies was as described for immunofluorescence, with centrifugation at 2200 rpm for 10 min at each step. Finally, cells were resuspended in 500 μL PI/RNAse solution for 30 min in the dark. Acquisition was performed using cytoflexS with a minimum of 1000 cells with the Cytexpert software. We excluded debris and doublets using morphometric parameters (FSC/SSC) and pulse area parameters from PI emission, respectively. To set the threshold of PAX6, TBR2, CTIP2, SATB2 and GFP-positive cells, control samples cells were incubated with secondary antibody and PI only. aRG progenitors DNA content was estimated by analyzing PI emission intensity of PAX6-positive cells only.

### Organotypic brain slice culture and drug treatment

Coronal slice of E13,5 control and CDC25B^cKO^ embryos were sectioned at 250 μm on a vibratome (Leica Microsystems). We deposited coronal slices on Millicell Cell Culture insert (0.4 μm, Millipore) in culture medium DMEM-F12, 1X penicilin/streptavidin. Slices remained incubated in 5% CO_2_ at 38°C for 24h. For drug treatment, brains slices were incubated in culture medium containing 1 μM PD0166285 (Selleckchem).

To determine cell proliferation and the PLM paradigm, 250 μM EdU was directly dropped off on each slice, for the duration required depending of the analysis (see above cell proliferation section). Brain slices were fixed 1h at room temperature in 4% paraformaldehyde, and, sectioned at 60 μm with a vibratome before immunostaining.

### DNA constructs and in utero electroporation

*In utero* electroporation experiments were performed in cortices of E13,5 embryos inside the pregnant mother as described in (Fawal et al. 2018). Embryos were harvested 28h later. Gain-of-function experiments were performed using a vector expressing two human Cdc25b forms hCDC25B3 (Cdc25b WT) or hCDC25B3DCDK (Cdc25bdelCDK) under the control of a cis regulatory element of the mouse Cdc25b called pccRE (Körner et al., 2001; (Bonnet et al. 2018). A control vector was generated with the ⍰ Gal gene downstream of the pccRE. 1,5 μg/μL CDC25B or control vector was co-injected with 1 μg/μL pCIG-GFP and Fast Green (Sigma) in the lateral ventricle of embryos neocortex, manually using a beveled and calibrated glass micropipette. For electroporation, five 50 ms pulses of 35 V with a 950 ms interval were delivered across the uterus with two 3-mm electrode paddles positioned on either side of the head. Before surgery, mice were subcutaneously injected with 100μL of morphine at 2.5 mg/kg. Mice were anesthetized with 4 % isoflurane. After surgery, mice were subcutaneously injected with 100μL of 2 mg/ml metacam.

### Imaging and statistical analysis

All the images of immunostained sections were acquired using a SP8 Leica confocal microscope. For each experiment, we analyzed at least three independent litters and three different slides per embryo. Quantitative data are expressed as mean ± s.e.m. Statistical analysis were performed using Graph Pad Prism 9 or the RStudio software. Significance was assessed by performing Mann-Whitney (MW) t-test, Wilcoxon test or using the mixed effect model (MEM) followed by an ANOVA test as described (Bonnet et al. 2018). * p < 0.05, ** p < 0.01, *** p < 0.001, ns. non significant.

## Acknowledgments

This work was supported by the Centre National de la Recherche Scientifique and Université P. Sabatier de Toulouse. Melanie roussat was funded by the Ministère de L’Enseignement Supérieur et de la Recherche (MESR). We thank Eric Agius and Valerie Lobjois for fruitful discussions. We thank Myriam Roussigné, Elise Cau, Mohamad Fawal for critical reading of the manuscript.

## SUPPLEMENTAL FIGURES

**Figure 2 - supplement 1:**
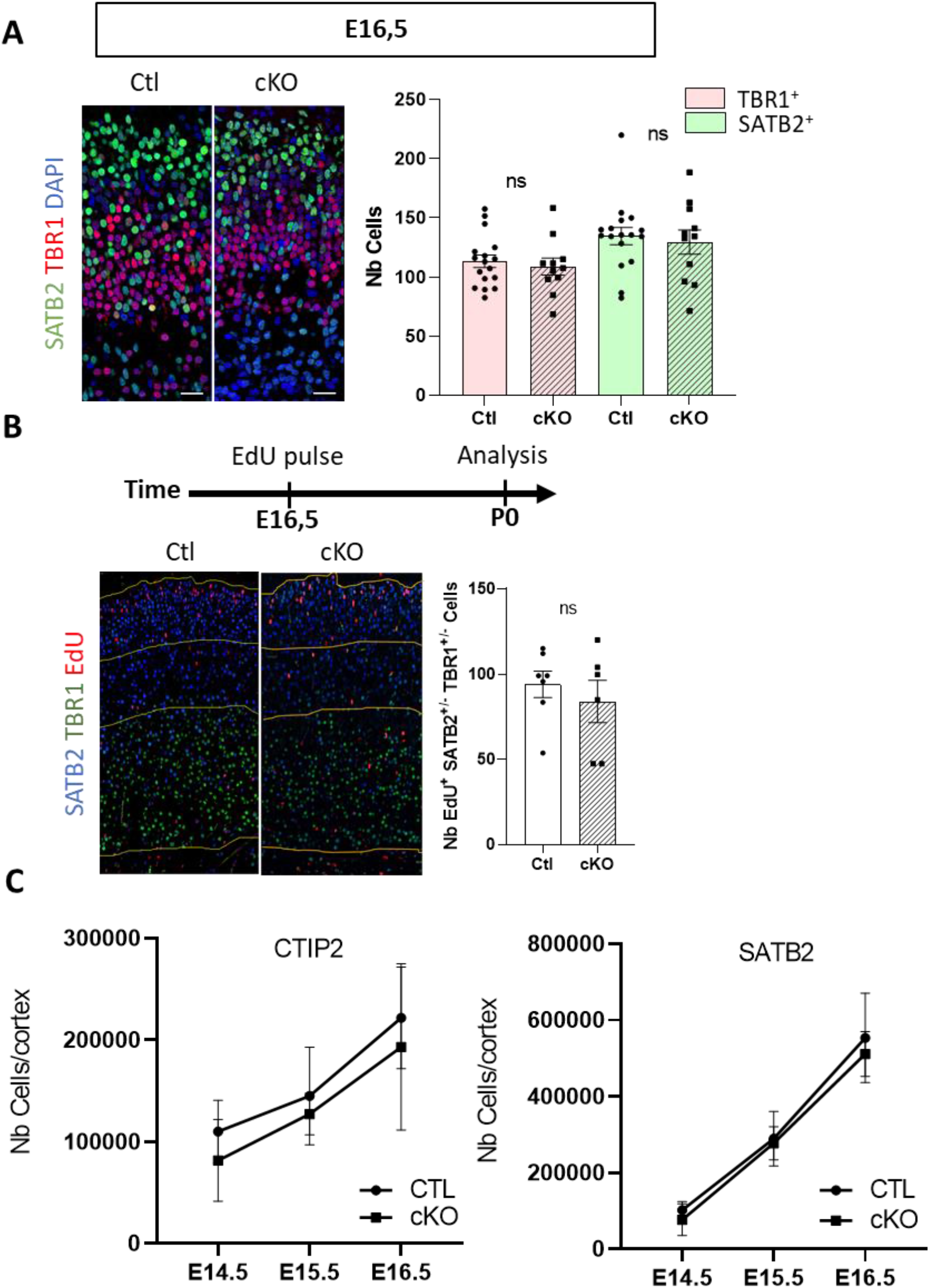
Neuronal production is progressively recovered in Cdc25b cKO embryos. **A**, SATB2+ and TBR1+ neurons quantification in Ctl and cKO embryos at E16.5. Each point is the mean value of 3 sections/embryo. Ctl n= 17, cKO n= 11. Mixed model. **B,** Birth dating experiment: quantification of EdU+ cells among SATB2+ and TBR1+ neurons at P0 in Ctl and cKO condition, following EdU pulse at E16.5. Each point represents the mean value of 3 sections/embryo. Ctl n= 7, cKO n=6, Mixed model. **C,** Dynamic of neuronal production in Ctl and cKO quantified by FACS sorting of Ctip2 and Satb2 neurons between E14.5 to E16.5. Each point represent the mean number of cells of several embryos: E14.S, Ctl n=6, cKO n=5; E1S.S, Ctl n= 6 cKO n=3 ; E16.5, Ctl n=6, cKO n=2. ns. Non significant. Scale bars represent 20 μm

**Figure 3 - supplement 1:**
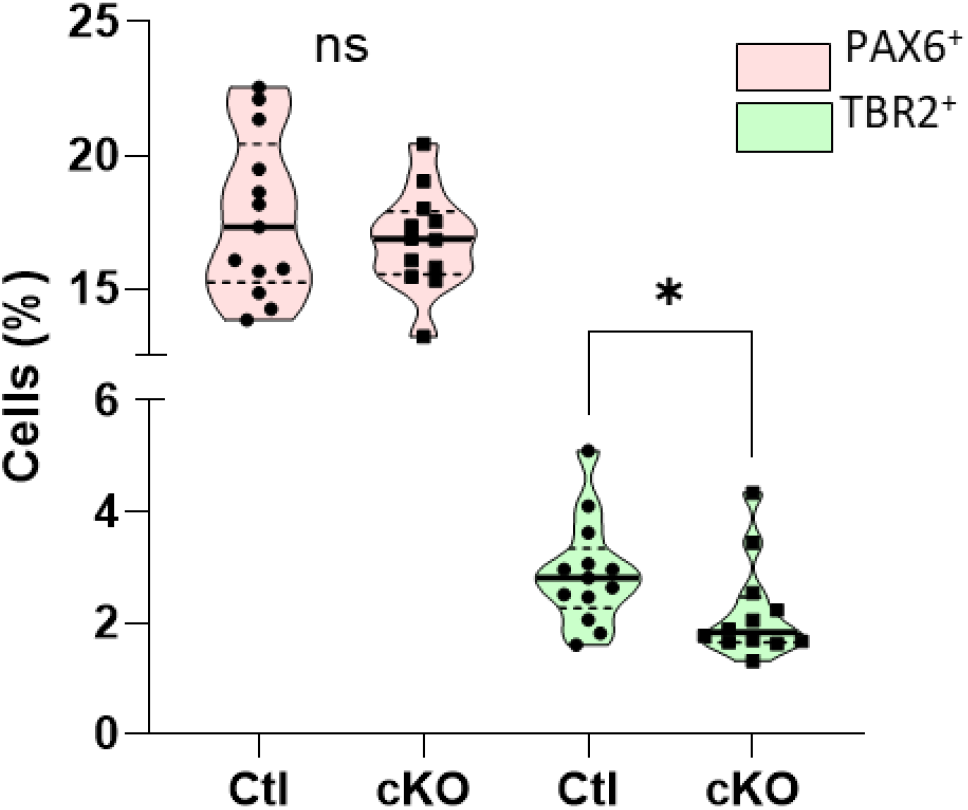
Validation of the reduction of TBR2 progenitors in cKO by FACS sorting. Number of PAX6+ (aRGs) and TBR2+ -blPs) progenitor cells in Ctl and cKO quantified by FACS sorting. The number of TBR2+ cells is significantly reduced in cKO. Flow cytometry analysis of the proportion of PAX6 + (aRGs + bRGs) and TBR2+ (blPs) singulates cells for Ctl and cKO. Each point represents the percentage of cells for an embryo. Ctl n=13, cKO n=12 Mann Whitney. * p<0,05

**Figure 3 - supplement 2:**
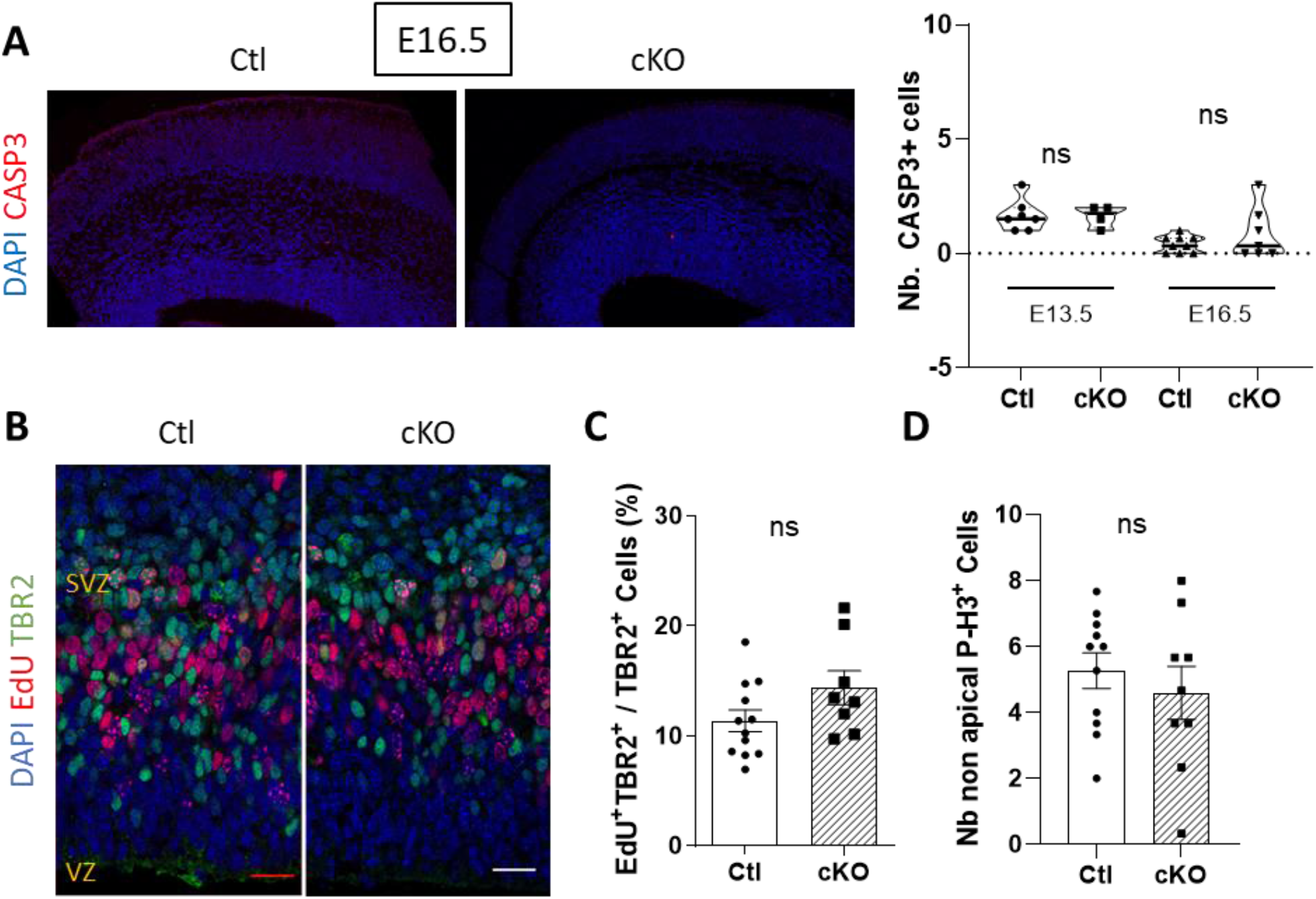
Reduction of blPs cells in the VZ is not due to cell death or reduced proliferation rate. **A**, Quantification of activated CASP3+ cells on frontal brain sections of Ctl and cKO embryos at E16.5. Each point is the mean value of counting in 3 hemispheres/embryo. E13.5: Ctl n=7, cKO n=4; E16.5: Ctl n=9, cKO n=7 Mann Whitney. **B,** Tbr2 and EdU labelling on coronal section after lH EdU pulse just before harvesting embryos at E14.5 **C,** Quantification of EdU+ cells among total Tbr2+ cells (blPs) in Ctl and cKO embryos at E14.5. Ctl n= 12, cKO n= 8. Mann Whitney. **D,** Quantification of P-H3+TBR2+ cells (non apical mitotic blPs) in E14.5 Ctl and cKO embryos. Ctl n= 11, cKO n= 9. Each point is the mean value of 3 sections/embryo. Mixed model. Ns, non significant. Scale bars represent 20 μm.

**Figure 3 - supplement 3:**
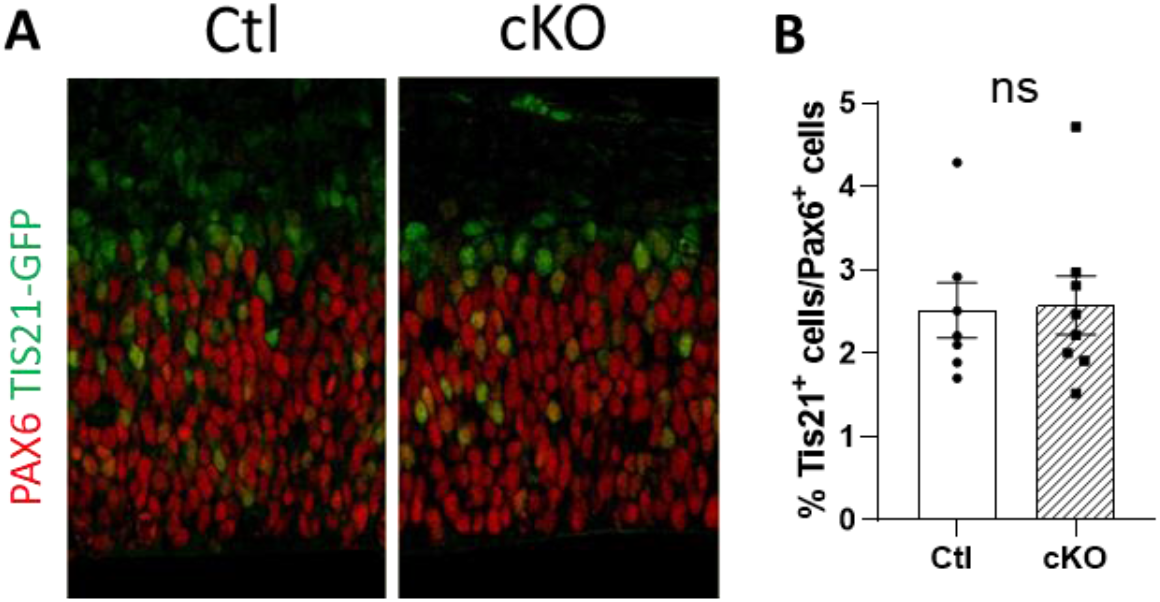
CDC25B does not change the division mode from proliferative to neurogenic in aRGs. **A**, PAX6 Immunostaining on brain coronal sections at E13.5 in TIS21-GFP+ Ctl or cKO embryos. **B,** Quantification of PAX6+ TIS21-GFP+ cells (differenciative aRGs) among PAX6+ cells. Each point is the mean value of 3 sections/embryo. Mixed model, ns, non significant. Scale bar is 20 μm.

**Figure 5 - supplement 1:**
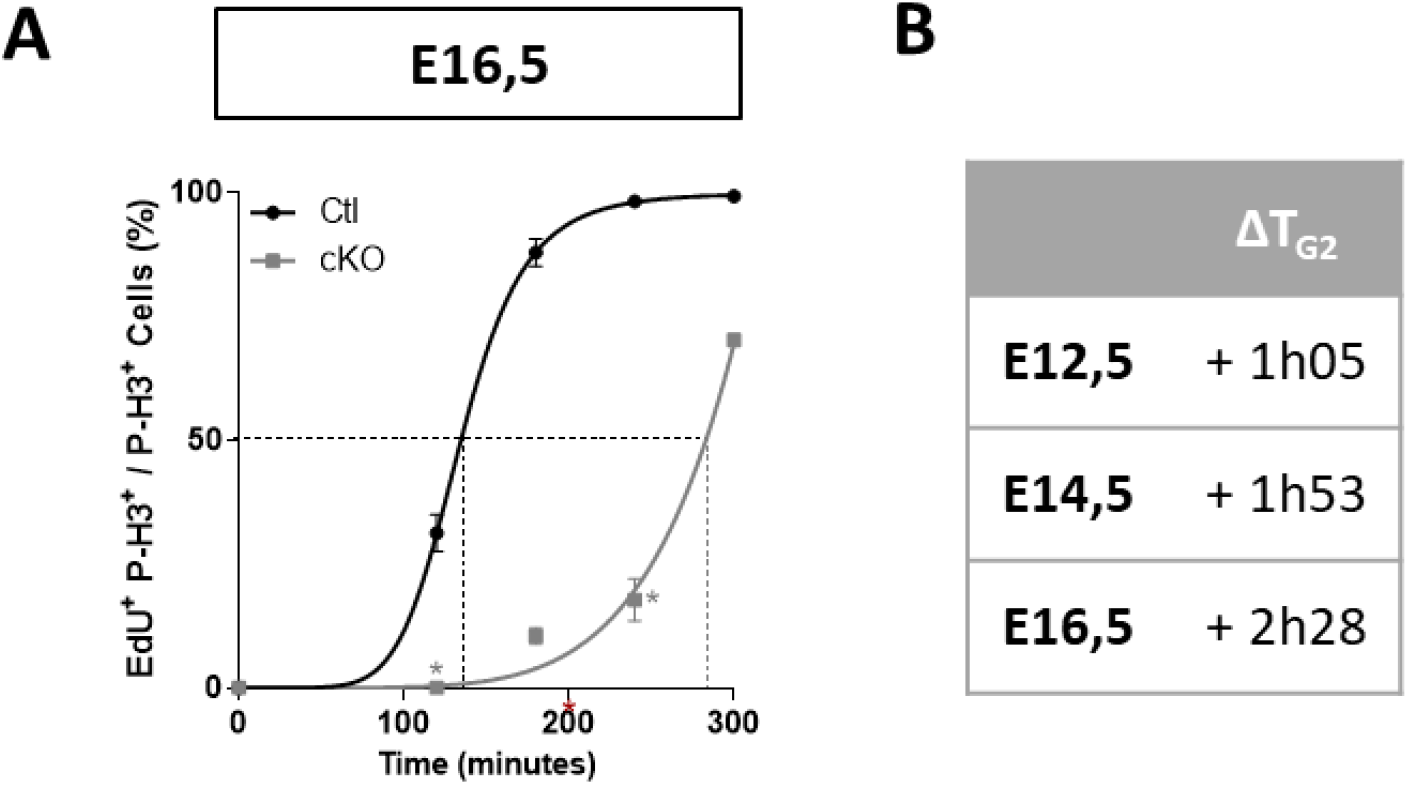
G2-phase length is still increased at E16.5 in cKO. **A**, Quantification of the proportion of apical (aRG) EdU^+^/P-H3^+^ among P-H3^+^ cells with increasing exposure at E16.5, in Ctl and cKO embryos. T_G2_ corresponds to the time when 50% of P-H3^+^ cells are EdU^+^ (indicated by the dotted lines). Mann-Whitney test. * p<0,05 **B,** T_G2_ difference between Ctl and cKO aRGs at E12.5, E14.5 and E16.5

**Figure 6 - supplement 1:**
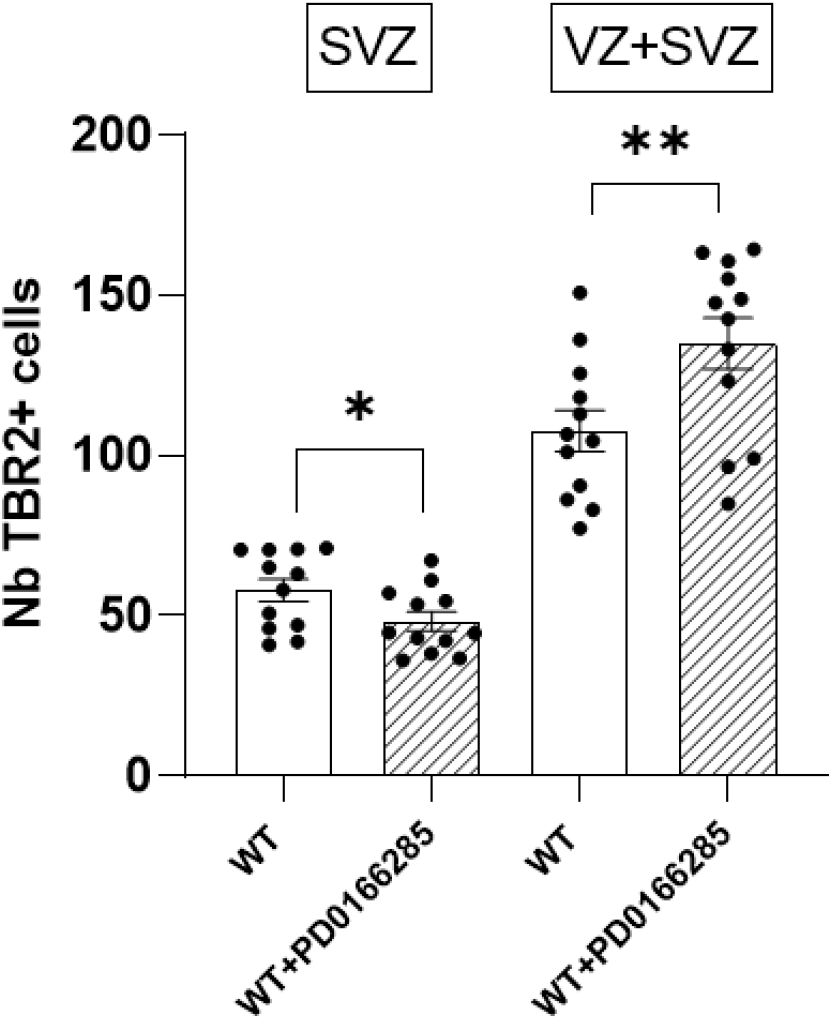
Modification of the number of TBR2+ cells (bIPs) following PD0166285 treatment in SVZ and VZ+SVZ. Quantification of the number of TBR2+ cells in the SVZ and VZ+SVZ in the same sections as in Figure 6F,G. (WT and WT+PD0166285 brain slices). Wilcoxon test. * p<0.05, **p<0.01. Each point represents the mean value of 3 sections/embryo. VZ = ventricular zone. Scale bars represent 20 μm.

